# The effect of counterfactual information on outcome value coding in medial prefrontal and cingulate cortex: from an absolute to a relative neural code

**DOI:** 10.1101/2020.01.08.898841

**Authors:** Doris Pischedda, Stefano Palminteri, Giorgio Coricelli

**Affiliations:** Center for Mind/Brain Sciences - CIMeC, University of Trento, 38123 Mattarello, Italy; Bernstein Center for Computational Neuroscience, Charité - Universitätsmedizin Berlin, 10115 Berlin, Germany; NeuroMI - Milan Center for Neuroscience, 20126 Milan, Italy; Human Reinforcement Learning team, Laboratoire de Neurosciences Cognitives et Computationnelles, Institut National de la Santé et de la Recherche Médicale, 75005 Paris, France; Département d’Études Cognitives, École Normale Supérieure, 75005 Paris, France.; Department of Economics, University of Southern California, 90089 Los Angeles, CA, USA.

## Abstract

Adaptive coding of stimuli in visual cortex is well documented in perception, where it supports efficient encoding over a broad range of possible percepts. Recently, a similar neural mechanism has been reported also in value-based decision, where it allows optimal encoding of vast ranges of values in PFC: neuronal response to value depends on the choice context (relative coding), rather than being invariant across contexts (absolute coding). Additionally, value learning is sensitive to the amount of feedback information: providing complete feedback (both obtained and forgone outcomes) instead of partial feedback (only obtained outcome) improves learning. However, it is unclear whether relative coding occurs in all PFC regions and how it is affected by feedback information. We systematically investigated univariate and multivariate feedback encoding in various PFC regions and compared three modes of neural coding: absolute, partially-adaptive and fully-adaptive.

Twenty-eight human participants (both sexes) performed a learning task while undergoing fMRI scanning. On each trial, they chose between two symbols associated with a certain outcome. Then, the decision outcome was revealed. Notably, in half of the trials participants received partial feedback, while in the other half they got complete feedback. We used univariate and multivariate analysis to explore value encoding in different feedback conditions.

We found that both obtained and forgone outcomes were encoded in mPFC, but with opposite sign in ventral and dorsal subdivisions. Moreover, we showed that increasing feedback information induced a switch from absolute to relative coding. Our results suggest that complete feedback information promotes context-dependent outcome encoding.

**SIGNIFICANCE STATEMENT:** This study offers a systematic investigation of the effect of the amount of feedback information (partial vs. complete) on univariate and multivariate outcome value encoding, within multiple regions in mPFC/ACC critical for value-based decisions and behavioural adaptation. Moreover, we provide the first comparison of three possible models of neural coding (i.e., absolute, partially-adaptive, and fully-adaptive coding) of value signal in mPFC by using commensurable measures of model accuracy. Taken together, our results help build a more comprehensive picture of how the human brain encodes and processes outcome value. In particular, our results suggest that simultaneous presentation of obtained and foregone outcomes promotes relative value representation.

## INTRODUCTION

Despite high variation in incoming information from the surrounding environment, humans perceive consistency in it. For example, an object can be seen in very different contexts (e.g., in daylight or in darkness), where its physical properties (e.g., color) vary greatly. Nonetheless, we perceive them as stable. To achieve this invariance, neurons adjust their sensitivity to the context characteristics through normalization, that is, they rescale their response to object properties relative to the specific context, instead of responding in absolute terms. These context effects allow efficient coding of broad ranges of sensory input and are well documented in perception (see, e.g., Carandini and Heeger, 2012; Louie and Glimcher, 2012). Recently, context-dependence and normalization were reported also in value-based decision, in both monkeys (Padoa-Schioppa, 2009; Bermudez and Schultz, 2010; Kobayashi et al., 2010; Rustichini et al., 2017; Conen and Padoa-Schioppa, 2019) and humans (Nieuwenhuis et al., 2005; Elliott et al., 2008; Cox and Kable, 2014; Burke et al., 2016; see Louie and De Martino, 2014 for a review), allowing optimal responses to vast ranges of values in PFC. Specifically, neurons seem to rescale their firing to adapt to the decision context (relative coding) so that their response to a specific value depends on the choice context (e.g., reward vs. punishment), rather than being invariant (absolute coding). Additionally, studies on value-based decision showed that feedback information affects value learning so that providing complete feedback (both obtained and foregone – counterfactual – choice outcomes) instead of partial feedback (only the obtained outcome) improves learning (Palminteri et al., 2015, 2017; Bavard et al., 2018). However, it is unclear (1) whether relative coding occurs in all PFC regions and (2) whether and how it is affected by feedback information.

Evidence for relative coding has been found in various PFC regions, such as mPFC, orbitofrontal cortex (OFC), and cingulate cortex (e.g., Nieuwenhuis et al., 2005; Elliott et al., 2008; Bunzeck et al., 2010; Cox and Kable, 2014), mainly with univariate fMRI analysis. This analysis identifies areas where neural response to value is consistent across voxels and participants. However, recordings from individual neurons showed that neuronal subpopulations in PFC exhibit opposite responses to value (e.g., Padoa-Schioppa and Assad, 2006; Schoenbaum et al., 2007; Kennerley and Wallis, 2009), thus questioning the ability of univariate analysis to capture all effects of interest. The introduction of multivariate methods (Haxby et al., 2001) that can detect value information encoded heterogeneously in brain activity patterns distributed across the brain, allowed to extend univariate results and answer questions related to the specific coding mechanisms the brain uses to encode value (Kahnt, 2018). Although with these techniques scientists could decode various value signals in the human brain (e.g., Clithero et al., 2009; Kahnt et al., 2010; Vickery et al., 2011; Wisniewski et al., 2015; Howard et al., 2016; Yan et al., 2016), the exact neural code that it uses to represent value is seldom investigated. The only decoding study in humans that investigated how outcome value adaptation occurs in the brain (Burke et al., 2016) considered only obtained outcomes. Moreover, despite the authors tested different possible types of adaptation in both univariate and multivariate signals, they employed methods that are hardly comparable.

To help build a more exhaustive picture, we designed an analysis plan to investigate outcome value processing and encoding in multiple regions in mPFC/OFC and cingulate cortex (see “Evaluating differences between output types”). The aim of the study is threefold: (1) to systematically evaluate univariate and multivariate effects in these regions, (2) to compare different coding models of outcome value encoding, and (3) to assess the effect of feedback information on value representation. We hypothesize that counterfactual information will produce rescaling of value signal depending on the context, such that the value of a neutral outcome becomes positive in a loss context (as absence of punishment) and negative in a gain context (as absence of reward), thus inducing relative coding of value signals.

## MATERIALS AND METHODS

### Participants

Twenty-eight participants took part in the experiment (16 females and 12 males; age 25.6 ± 5.4 years). All participants were right-handed, did not report any psychiatric or neurological history, and had normal or corrected-to-normal vision. The ethics committee of the University of Trento approved the study and participants provided written informed consent before their inclusion in the study. They received monetary payment, calculated as a show-up fee plus the total amount of money they won during the experiment.

### Experimental design, stimuli, and procedure

Participants performed an instrumental learning task while undergoing fMRI scanning. At the beginning of the experimental session, they received written instructions and the task was further explained orally, if necessary. Participants were requested to maximize their payoff, considering that both reward seeking and punishment avoidance were equally important strategies and that only factual (but not counterfactual, see below) outcomes would be used to calculate their total earnings. After the instruction phase, participants performed a training session to practice the task before entering the fMRI scanner. After practice, they performed four learning runs in the scanner. Participants were presented with a pair of abstract symbols belonging to the Agathodaimon alphabet. For each run, we used eight different symbols arranged in four pairs to produce four choice contexts (i.e., reward/partial, reward/complete, punishment/partial, and punishment/complete). The contexts were defined based on the possible outcome (either reward - winning 0.5€ vs. 0€ - or punishment - losing 0.5€ vs. 0€) and the feedback provided (either partial - only the outcome of the chosen option - or complete - the outcomes of both the chosen and the unchosen options). The contexts were characterized by a fixed pair of symbols within the same run but the pairs associated with each context differed among runs. Each symbol in one pair was associated with the possible outcomes with complementary probability (0.75/0.25 for reward and 0.25/0.75 for punishment). Participants performed 96 trials (24 repetitions of the four experimental conditions) during each scanning run.

Examples of trials with either partial (top rows) or complete (bottom rows) feedback information are shown in Figure 1. At the beginning of each trial, a pair of symbols was shown (with each symbol randomly presented either on the left or on the right of a central fixation cross). Participants had to choose one of the symbols and to press the corresponding button with their left or right thumb, within 3,000ms. Afterward, a red pointer was displayed under the selected option for 500ms, which was then replaced by its outcome (+0.5€, 0.0€, or −0.5€) shown for 3,000ms. In complete feedback trials, the outcome of the unchosen option (i.e., counterfactual) was displayed as well. The next trial followed after about 1,000ms (jittered, minimum 500ms and maximum 1,500ms) during which a fixation screen was shown. Presentation order of the different pairs of symbols and the position of each symbol in the pair was pseudo-randomized and unpredictable so that each symbol was displayed an equal number of times to either side of the screen.

**Figure 1.**
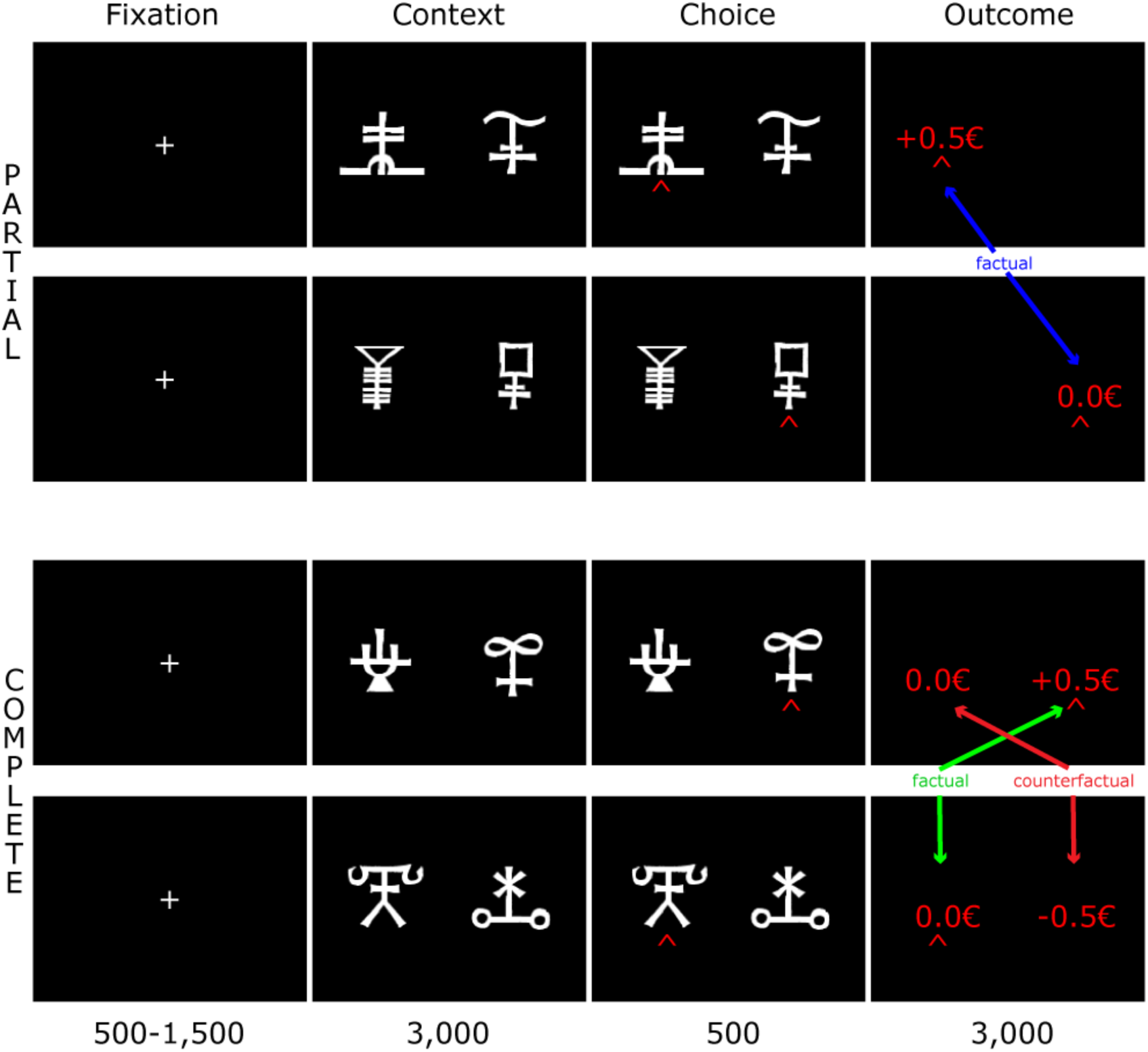
Experimental paradigm. Examples of experimental trials with either partial (top rows) or complete (bottom rows) feedback. Outcome types (factual vs. counterfactual) are also specified. From these two factors (i.e., feedback and outcome type), three different experimental conditions originate: partial factual, complete factual, and complete counterfactual outcomes. Adapted from Palminteri et al. (2015 p.3).

At the end of the four experimental runs and while the anatomical image was acquired, participants performed a post-learning evaluation of option value. We do not describe this task here since the data were not used for the analyses described in this paper. For details, please see the original article (Palminteri et al., 2015).

### Image acquisition and preprocessing

fMRI images were recorded using a 4T Bruker MedSpec Biospin MR scanner (CIMeC, Trento, Italy) equipped with an eight-channel head coil. For each of the 4 scanning runs, we collected 338 T2*-weighted EPI volumes. Each image consisted of 47 slices recorded in ascending interleaved order with acquisition parameters as follows: TR 2,200ms, TE 21ms, FA 75°, FOV 192mm, matrix size 64 × 64, yielding an in-plane voxel resolution of 3mm^3^ with voxel size of 3mm × 3mm × 2.3mm. A T1-weighted anatomical sequence was recorded as well, with imaging parameters: TR 2,700ms, TE 4.18ms, FA 7°, matrix size 224 × 256 × 176, with a voxel resolution of 1mm^3^.

We used SPM12 (RRID:SCR_007037) to pre-process and analyze the fMRI data. During pre-processing, the images were realigned and slice-time corrected, low-frequency noise was removed using a high-pass filter with a cutoff period of 128s (Worsley and Friston, 1995), and an autoregressive AR model was fitted to the residuals to allow for temporal autocorrelations (Friston et al., 2002). The anatomical image was segmented using the template tissue probability maps in SPM12 and used as a reference to coregister the functional images. To perform the univariate analyses, fMRI images were also smoothed (FWHM 6mm) and normalized to MNI space. For the multi-voxel pattern analysis (MVPA), the images were neither normalized nor smoothed to preserve fine-grained patterns of brain activity.

### Statistical analyses

fMRI data from this experiment has been previously analyzed with a model-based procedure to test the hypothesis that successful avoidance of punishment is reframed as a positive outcome and that neural activity in brain regions within the valuation system is better accounted by a relative model of value representation as compared with an absolute model (Palminteri et al., 2015). An additional aim of the previous analyses was to assess neural encoding of choice and outcome in those different regions as a function of task context.

The analyses described in this manuscript differ in either the method used (MVPA) or the question asked (whether and how outcome representation changes as a function of the available information).

#### Behavioral analyses

For the sake of clarity and completeness, we report behavioral results (see Figure 2) from the original paper (Palminteri et al., 2015). We used one sample t-tests to assess learning in the different experimental conditions (i.e., to compare the actual correct choice rate with the value expected by chance). To assess possible effects of context (i.e., reward vs. punishment) and feedback information (i.e., partial vs. complete) on either accuracy or reaction times (RT), we performed linear mixed-effects model (LMM) analysis with accuracy (or RT) as predicted value and context and feedback information as fixed effects. As random effect, we introduced intercepts for each participant, thus allowing inter-subject variability in behavioral responses.

**Figure 2.**
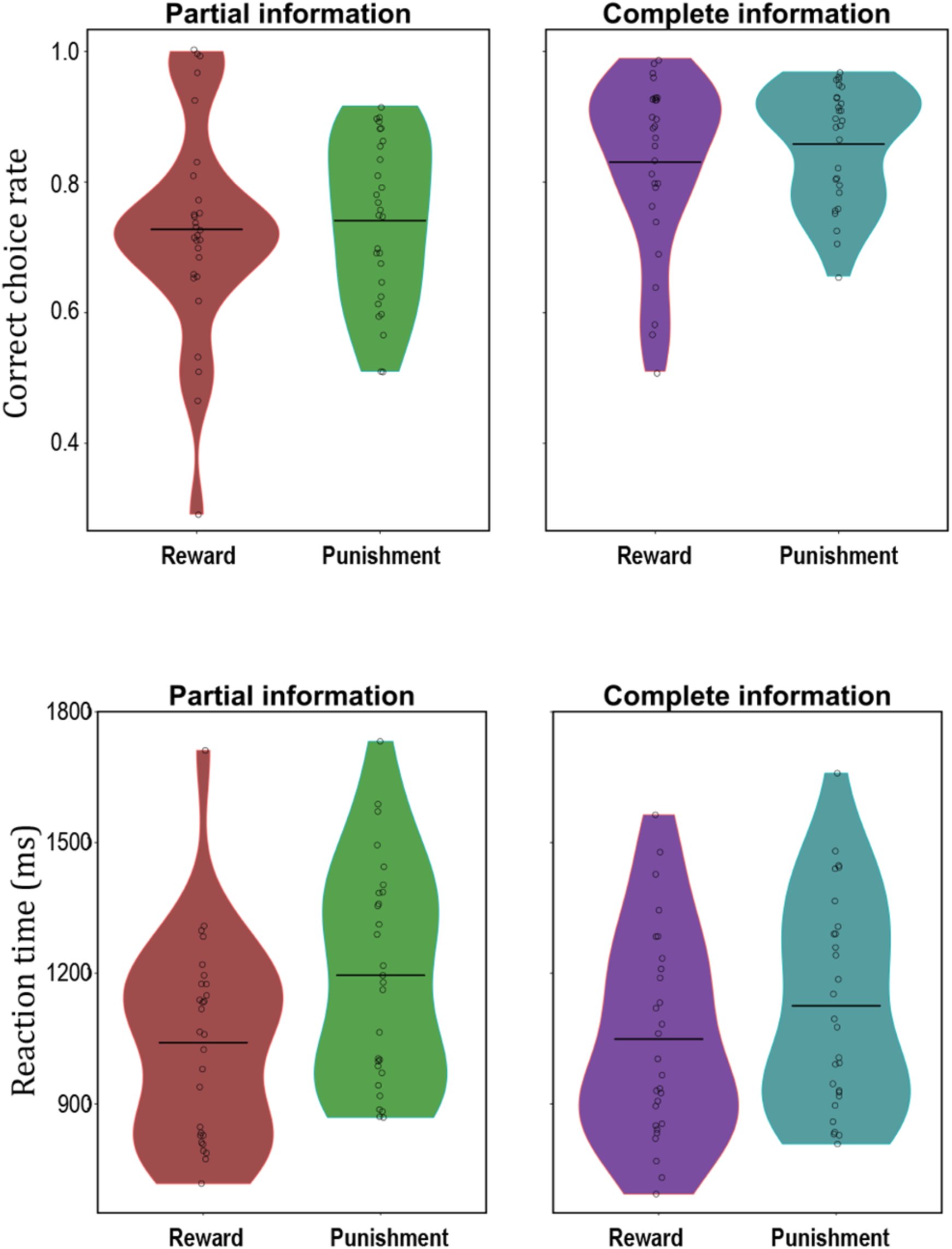
Behavioral results. The graphs on the top row display the correct choice rate in the partial (left) and complete (right) information conditions for the two contexts. In the bottom row, mean reaction times in partial (left) and complete (right) information trials are depicted.

### Evaluating differences between output types

#### ROI selection

The first aim of the present study was to assess univariate and multivariate effects of outcome encoding in mPFC and cingulate cortex and to identify possible differences between distinct outcome conditions (see “Experimental design, stimuli, and procedure”). In particular, we aimed to test whether the regions along the dorsal/ventral axis of mPFC were sensitive to outcome processing or encoded information about outcome value for the three outcome types we investigated. To avoid circular inference (Kriegeskorte et al., 2009), we selected an independent set of ROIs from the Brainnetome Atlas (Fan et al., 2016; RRID:SCR_014091). We included nine areas along mPFC/OFC and cingulate cortex in the set of selected ROIs (see Figure 3A), since these areas have been previously implicated in reward processing, especially when choice outcome is revealed (Knutson et al., 2003; Diekhof et al., 2012; Clithero and Rangel, 2014) as compared to, for example, ventral striatum, which seems to be more strongly (or equally) activated during outcome anticipation (Knutson et al., 2001; Rangel et al., 2008; Diekhof et al., 2012; Stott and Redish, 2014; Oldham et al., 2018) and to reflect prediction error rather than value (Hare et al., 2008; Rohe et al., 2012).

**Figure 3.**
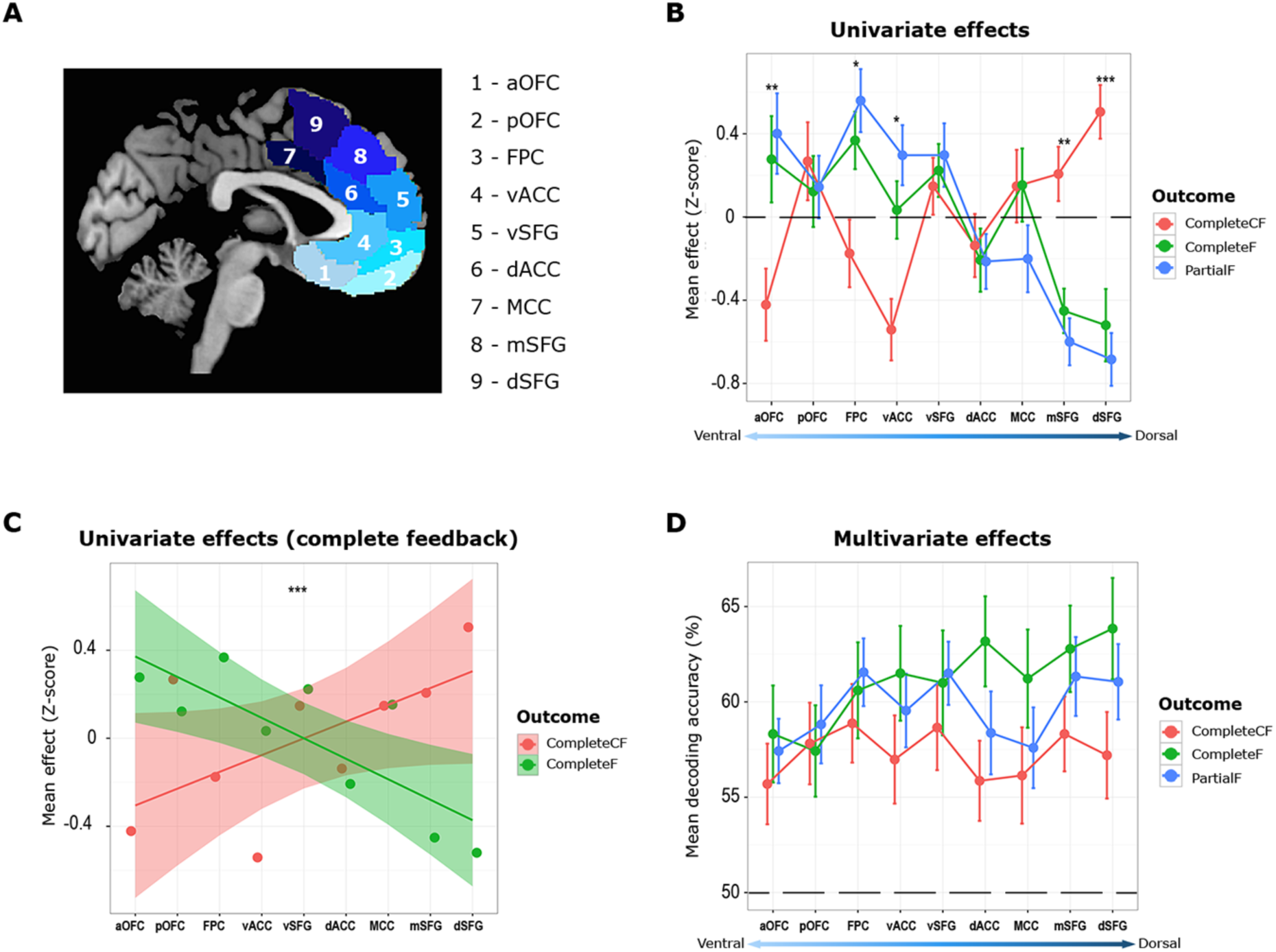
Univariate and multivariate effects in the selected ROIs. (A) The selected ROIs are depicted. fMRI analyses were conducted on nine preselected anatomical ROIs along the ventral/dorsal axis of mPFC. The set of ROIs comprises one dorsal, one middle, and one ventral area for each of three target regions: superior frontal gyrus, orbitofrontal/frontopolar cortex, and cingulate cortex. aOFC = anterior orbitofrontal cortex; pOFC = posterior orbitofrontal cortex; FPC = frontopolar cortex; vACC = ventral anterior cingulate cortex; dACC = dorsal anterior cingulate cortex; MCC = middle cingulate cortex; vSFG = ventral SFG; mSFG = middle SFG; dSFG = dorsal SFG. (B) Results of the univariate analyses on outcome valence are shown. ROIs on the x-axis are arranged from the most ventral (left) to the most dorsal (right), according to their mean z-coordinate. The dashed line indicates a null effect. ROIs where there was a significant difference between counterfactual outcomes and both factual outcomes are highlighted. (C) Univariate effects are shown only for outcomes in trials with complete feedback. The order of the ROIs on the x-axis is as in (B). The interaction effect outcome type x ROI was significant, reflecting positive encoding of value for factual outcomes and negative encoding for counterfactual outcomes in more anterior mPFC ROIs but the opposite pattern in the dorsal regions. (D) The graph displays the results of the multivariate analyses on outcome valence. The dashed line marks chance level (rounded-up to 50, actual chance level calculated with shuffle tests is 49.997). ROIs on the x-axis are arranged as in (B) and (C). No ROI is highlighted on this graph as neither the main effect of ROI nor the interaction effect outcome type x ROI was significant in this analysis. Error bars display SEM. *** = *p* < .001; ** = *p* < .01; * = *p* < .05.

#### Univariate analyses

Realigned, slice-time corrected, smoothed, and normalized images were used to obtain parameter estimates for the GLM. The GLM was set up to assess processing and encoding of outcome value in trials with either partial or complete information (see “Experimental design, stimuli, and procedure” for details on the different information conditions). Twelve regressors were included in the model, corresponding to the four possible outcomes (i.e., +0.5€, +0.0€, -0.0€, and −0.5€) for each of the three outcome types (i.e., partial factual, complete factual, and complete counterfactual). We defined the time vectors for the twelve regressors using the onset time of the outcome screen presentation; all regressors were convolved with the canonical HRF function to compute the GLM. The estimates were linearly combined to contrast the conditions of interest: estimates for negative outcomes (i.e., +0.0€ and - 0.5€) were subtracted from those for positive outcomes (i.e., +0.5€ and -0.0€) for each outcome type.

#### Multivariate analyses

Realigned and slice-time corrected images were used to estimate run-wise correlation coefficients for the GLM (see above). We employed MVPA with an ROI-based approach to determine which brain regions contained information about outcome value. Three decoding analyses were performed: the purpose of the first one was to identify brain regions encoding information about outcome value in trials with partial feedback, the second one aimed at identifying brain regions containing information about factual outcome value in trials with complete feedback, and the goal of the last analysis was to establish whether any region in the brain contained information about the counterfactual outcome. All analyses decoded between positive and negative outcomes (i.e., +0.5€/-0.0€ vs. +0.0€/-0.5€, see “Experimental design, stimuli, and procedure”) using the parameter estimates from the GLM (see above). Decoding analyses tested whether patterns of neural activity within each ROI contained information allowing for distinguishing between two task conditions (e.g., a positive and a negative outcome). We performed pairwise decoding between positive and negative outcomes. For each selected ROI and for each of the four fMRI runs, we extracted parameter estimates for the four experimental conditions (i.e., for +0.5€, +0.0€, -0.0€, and −0.5€ outcomes) for the outcome type of interest. A total of 16 pattern vectors (one for each of the 4 conditions and for each of the 4 runs) were available for each participant. For each decoding analysis, the pattern vectors of the pair under evaluation were assigned to independent training and test sets, to avoid overfitting (Duda et al., 2000). We implemented a leave-one-run-out cross-validation procedure, where data from each run were assigned, in turn, to the test set and data from the remaining three runs were used to train a support vector classifier (Müller et al., 2001; Cox and Savoy, 2003; here, regularization parameter C = 1) to distinguish between the two conditions of interest. The trained classifier was applied to data from the test set and its classification accuracy was calculated as the ratio between the number of correct classifications and the number of all classifications that were performed. The resulting classification accuracy reflects how well the classifier distinguished between the two conditions of interest in the specified ROI. For each of the three analyses, the decoding procedure was repeated using all possible condition pairs as classes for the classification. Accuracies resulting from each pairwise classification were averaged, yielding an accuracy value for each ROI, participant, and analysis. We used shuffle tests with all possible permutations of the condition labels to calculate the actual chance level. To assess statistical significance, we randomly sampled from the results of the permutations and calculated second level statistics by comparing actual accuracies with the distribution of accuracies obtained from 10^6^ resamplings. All decoding analyses were performed using The Decoding Toolbox (TDT Hebart et al., 2015; RRID:SCR_017424).

#### ROI analyses

We employed ROI analyses to assess the effect of different types of outcomes at the neural level in the set of selected ROIs. For each participant and for each analysis (see above), we used the mean contrast effect (or decoding accuracy) for each ROI. Then, we used LMMs to assess the effect of outcome type and ROI on the extracted mean estimates (see “Univariate and multivariate effects of outcome valence” for details on the single analyses). We ran LMM analyses using R 3.2.3 (RRID:SCR_001905) and the lme4 package (Bates et al., 2015; RRID:SCR_015654). We favored LMMs over repeated-measures ANOVA because they take into account subjects’ sensitivity to different experimental conditions beyond individual variability in mean responses. Thus, LMMs are more powerful than repeated-measures ANOVA (Barr et al., 2013). Statistical significance of the tests was calculated with likelihood ratio tests of the model with the effect of interest against the model without that effect (Barr et al., 2013). Post hoc tests comparing different levels of the factors and interactions showing a significant effect were performed using the emmeans package (Lenth, 2018). To maintain the alpha level at the intended value of 0.05, we performed pairwise comparisons with the Tukey HSD test (Tukey, 1949) and chose the Holm’s method (Holm, 1979) to correct for multiple comparisons and control the family-wise error rate while maintaining high statistical power.

### Comparing different value coding models

The second goal of this study was to assess how the human brain encodes outcome value. Specifically, we aimed to test whether outcomes are represented using an absolute, a partially-adaptive, or a fully-adaptive code (see Figure 4A). To this purpose, we implemented different support vector regression (SVR) analyses (see Kahnt et al., 2011 for a detailed description) that captured the characteristics of each of the theoretical models. Different from the classification approach described in “Evaluating differences between output types”, we used the local patterns of brain activity associated with multiple predictors (i.e., the four possible outcomes: +0.5€, +0.0€, -0.0€, and −0.5€) to predict a continuous variable (i.e., the outcome value as assumed by the theoretical model). We adopted a leave-one-run-out cross-validation procedure (see “Evaluating differences between output types”) to assure independence between train and test sets (Mitchell, 1997). For each ROI, we calculated the prediction accuracy of the SVR model using the Fisher’s Z-transformed correlation coefficient between the predicted outcome values and the actual values in the test set. The SVR analyses were implemented using TDT (Hebart et al., 2015; RRID:SCR_017424).

**Figure 4.**
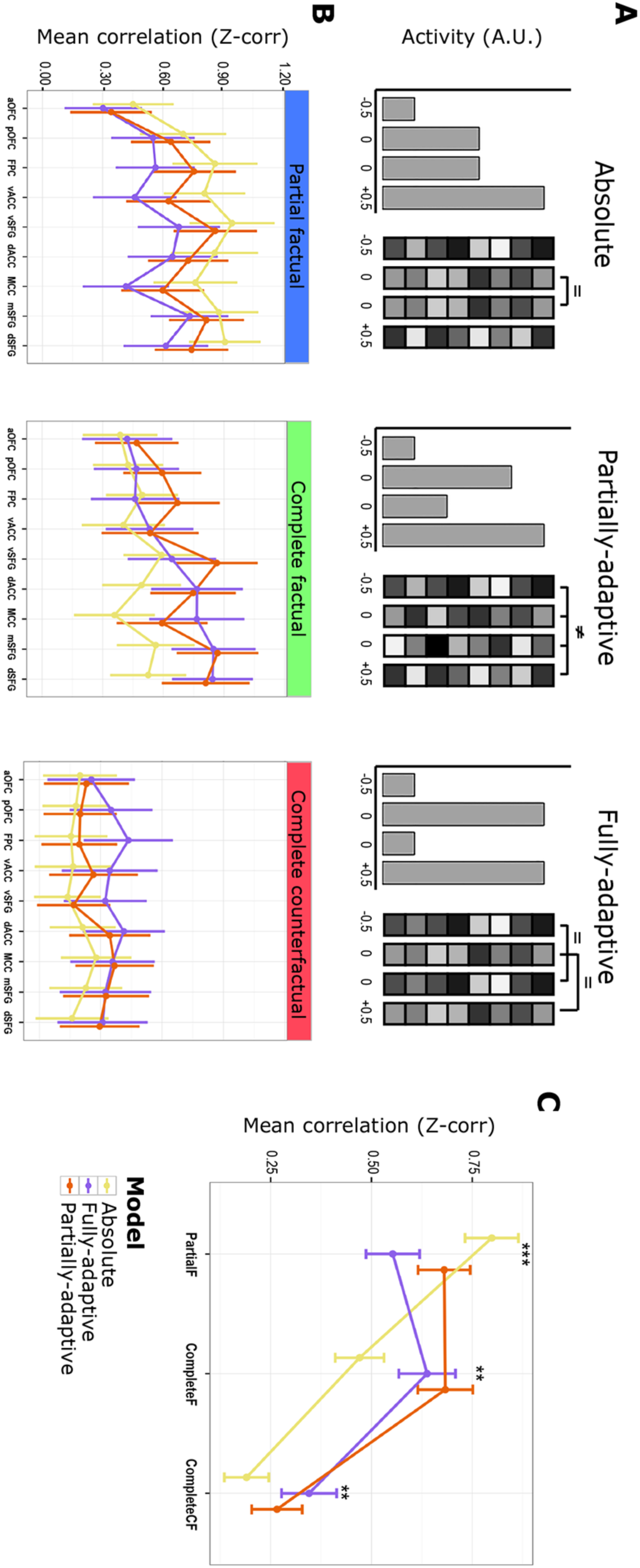
Comparison of different value coding models. (A) Three different models for value coding are shown. Under absolute value coding (left panel), values are represented independent of the context, thus neural activity patterns encoding the same value in reward and punishment contexts (e.g., +0.0€ and - 0.0€) are equivalent. With partially-adaptive coding (middle panel), values are rescaled so that positive outcomes in punishment contexts (e.g., -0.0€) have a higher value than negative ones in reward contexts (e.g., +0.0€). Finally, if the code is fully adaptive (right panel) neural patterns encoding a positive outcome in different contexts are equivalent, but different from those encoding negative outcomes, which are identical as well. (B) Results of the analysis comparing different value coding models. The absolute coding model is the best at predicting neural activity patterns representing factual outcome when partial feedback is provided (left panel) in all ROIs. Instead, for counterfactual outcome value encoding (right panel) the fully-adaptive coding model is the one with higher prediction accuracy (see “Comparing different value coding models”). For factual outcome encoding when complete feedback is given (middle panel), the model with higher prediction accuracy is either the fully-adaptive or the partially-adaptive one, depending on the ROI. (C) Interaction effect of coding model x type of outcome. The absolute model had higher prediction accuracy for factual outcomes with partial information than both the partially-adaptive and the fully-adaptive models. Instead, the fully-adaptive model had higher prediction accuracy than the absolute model for factual and counterfactual outcomes with complete feedback. aOFC = anterior orbitofrontal cortex; pOFC = posterior orbitofrontal cortex; FPC = frontopolar cortex; vACC = ventral anterior cingulate cortex; dACC = dorsal anterior cingulate cortex; MCC = middle cingulate cortex; vSFG = ventral SFG; mSFG = middle SFG; dSFG = dorsal SFG. Error bars represent SEM. *** = *p* < .001; ** = *p* < .01.

The first analysis aimed to identify brain regions representing value with an absolute code. Under absolute value coding, neural patterns of brain activity should reflect the magnitude of the reward, irrespective of the context (see Figure 4A, left panel). Thus, local pattern of brain activity encoding neutral outcomes in the reward (i.e., +0.0€) and in the punishment (i.e., -0.0€) contexts should be equivalent; in addition, they should be different from neural patterns encoding both the best (i.e., +0.5€) and the worst (i.e., −0.5€) outcome. To identify brain regions showing neural activity patterns in line with this model, we used a SVR model where we set three different outcome values: one value (i.e., 0) for both the neutral outcomes, one value (i.e., 2) for the best outcome, and one value (i.e., -2) for the worst outcome.

In the second analysis, we tested whether any brain regions encoded outcome value in a fully-adaptive way. If a region encoded outcomes with a fully-adaptive code, good and bad outcomes should be represented in the same way, irrespective of the context in which they occur (see Figure 4A, right panel). Thus, neural patterns encoding the good outcome in the reward context (i.e., +0.5€) and in the punishment context (i.e., -0.0€) should be equivalent; similarly, brain activity patterns representing the bad outcome in the punishment context (i.e., −0.5€) and in the reward context (i.e., +0.0€) should be identical. To assess whether neural activity patterns encoded outcome value according to the fully-adaptive coding model, we performed SVR assigning the same outcome value (i.e., 2) to both the good outcomes (i.e., +0.5€ and -0.0€) and a different value (i.e., -2) to both the bad outcomes (i.e., +0.0€ and −0.5€).

Finally, a third analysis assessed partially-adaptive coding of outcome value. If the human brain represented outcomes with a code that is just partially-adaptive, the value of the good outcome in the punishment context (i.e., -0.0€) should not be as high as the value of the good outcome in the reward context (i.e., +0.5€) and the value of the bad outcome in the reward context (i.e., +0.0€) should not be as low as the value of the bad outcome in the punishment context (i.e., −0.5€). However, the value attributed to the null outcome (i.e., 0.0€) should be higher in the punishment context as compared to the reward context, since the absence of punishment is a more positive outcome than the absence of reward, as expected by the partially-adaptive coding model (see Figure 4A, middle panel). To test whether local patterns of brain activity reflected a partially-adaptive coding, we set a SVR model in which different values were assigned to each of the possible outcomes (i.e., 2 and 1 to the good outcomes and -1 and -2 to the bad outcomes).

The three SVR analyses were performed on data from each pre-selected ROI (see “Evaluating differences between output types”), yielding to a prediction accuracy value for each participant, model, analysis, and ROI. We used LMM and likelihood ratio tests (see “Evaluating differences between output types”) to assess the effect of coding model, outcome type, and ROI on prediction accuracy (see “Revealing the neural code underlying outcome valence representation” for details of each analysis).

### Code Accessibility

All analyses were performed using in-house developed code and implemented either in MATLAB (RRID:SCR_001622) or R (RRID:SCR_001905). All code and the data for the behavioral and the ROI analyses are available through GitLab: https://gitlab.com/doris.pischedda/value_normalization.git.

## RESULTS

### Behavioral results

An extensive description of the behavioral results from this experiment can be found in the original paper (Palminteri et al., 2015). Here, we report only a summary of them, together with the findings of the neuroimaging analyses described in the previous section (see “Statistical analyses”), to provide a full picture of the results of the experiment.

Mean accuracy and RT in the different experimental conditions are shown in Figure 2. Behavioral data provide evidence for learning, as the percentage of correct responses for participants was significantly higher than what would be expected by chance (i.e., 50%) in all contexts (all *T*s > 7.4, all *p*s < 6 x 10^-8^). Linear mixed-effects model (LMM) analyses (see “Statistical analyses” for details) showed a main effect of feedback information on accuracy (*χ^2^*(1) = 34.4, *p* = 3 x 10^-8^) but not on RT (*p* = .08) and a main effect of context and an interaction effect of context and information on RT (*χ^2^*(1) = 34.8, *p* = 4 x 10^-9^ and *χ^2^*(1) = 5.3, *p* = .021, respectively), but not on accuracy (*p* = .35 and *p* = .70, respectively). Post hoc test showed higher accuracy when feedback information was complete as compared to when it was partial (*p* < .0001) and slower RT when partial rather than complete feedback was provided (*p* = .004), but only in punishment trials. Overall, the behavioral results suggest that people learn similarly in reward and punishment contexts and can integrate counterfactual information to improve their performance.

### Univariate and multivariate effects of outcome valence

The first aim of the analyses described in this manuscript was to investigate *where* the human brain represents information about outcome value during a learning task including both reward and punishment contexts and where either partial or complete feedback was provided. In particular, we explored both univariate and multivariate effects of outcome type in a set of preselected regions along the dorsal/ventral axis of mPFC to confirm their role in processing/encoding of outcome value. As our analyses rely on both differences between positive and negative outcomes and monetary gain and losses (see Statistical analyses), we describe our results in terms of outcome valence rather than value (see, e.g., Kahnt et al., 2014 for extensive discussion).

To assess possible differences in univariate effects between different outcome types and ROIs, we performed an LMM analysis modeling brain signal change as predicted value and outcome type and ROI as fixed effects. We considered intercepts for each participant as random effect, thus allowing inter-subject variability in mean brain activation. Results are shown in Figure 3B. Both the ROI and the outcome type x ROI interaction had a significant effect on signal change (*χ^2^*(8) = 36.7, *p* = 1.3 x 10^-5^ and *χ^2^*(16) = 96.8, *p* = 1.4 x 10^-13^, respectively). Instead, the main effect of outcome type on the neural signal was not significant (*χ^2^*(2) = 0, *p* = 1). Post hoc tests performed using the Tukey HSD test (corrected for multiple comparisons with the Holm’s method) revealed higher univariate effects for factual than for counterfactual outcomes (all *p*s < .026) and no difference for factual outcomes between the partial and the complete feedback condition (all *p*s > .22) in all regions, except in middle (mSFG) and dorsal (dSFG) superior frontal gyrus, where univariate effects were higher for counterfactual than for factual outcomes (all *p*s < .006), and in posterior OFC (pOFC), ventral superior frontal gyrus (vSFG), in dorsal ACC (dACC), and middle cingulate cortex (MCC) where there was no difference between the three outcome conditions (all *p*s > .31). Taken together, the univariate results indicate that, while anterior mPFC regions (except pOFC) encode factual outcomes (either with partial or complete information) positively and counterfactual outcomes negatively, the opposite is true for the most posterior areas (see Figure 3C), suggesting that ventral regions process primarily factual outcome information while more posterior areas play a main role in counterfactual outcome processing.

To evaluate possible differences in multivariate effects between different ROIs and outcome types, we used an LMM modeling decoding accuracy as predicted value and ROI and type of outcome as fixed effects. As random effect, we added random intercepts for participants to account for individual differences in participants’ mean neural response. Results are shown in Figure 3D. The effect of outcome type on decoding accuracy was significant (*χ^2^*(2) = 17.4, *p* = .0002), reflecting better decoding for factual outcome as compared to counterfactual ones. Instead, neither the effect of ROI nor the outcome type x ROI interaction reached significance (*χ^2^*(8) = 10.9, *p* = .21 and *χ^2^*(16) = 8.1, *p* = .95, respectively). Post hoc tests (with Tukey HSD, corrected for multiple comparisons with the Holm’s method) showed worse decoding for counterfactual outcomes than for factual outcomes with either complete (*p* = .0001) or partial (*p* = .019) feedback and no difference in factual outcome encoding between the partial and complete feedback conditions (*p* = .13). In summary, multivariate results show that information about counterfactual outcomes can be decoded in multiple regions of mPFC, although with lower accuracy than for factual ones. As counterfactual outcomes are not relevant for the obtained reward and representational strength is stronger for attended than unattended items (e.g., Christophel et al., 2018), this suggests that they receive less attention.

### Revealing the neural code underlying outcome valence representation

The main goal of this study was to investigate *how* the human brain represents outcome information when the result of the chosen (and unchosen) option is revealed. To this purpose, we compared three different models of valence coding (i.e., absolute, partially-adaptive, and fully-adaptive models, see Figure 4A) to assess which one would account better for valence representation of the three possible outcomes (i.e., factual outcome with partial or complete feedback and counterfactual outcome) in the different ROIs.

To test for possible differences in prediction accuracy (see “Comparing different value coding models”) among the three coding models for each outcome type and in the different ROIs, we used an LMM modeling prediction accuracy as the dependent measure and coding model, outcome type, and ROI as fixed effects. As random effect, we included intercepts for each participant (see “Univariate and multivariate effects of outcome valence”) in the model. Mean prediction accuracies for the different conditions are shown in Figure 4B. The LMM analysis showed a main effect of both outcome type (*χ^2^*(2) = 227.0, *p* < 2.2 x 10^-16^) and ROI (*χ^2^*(8) = 52.1, *p* = 1.6 x 10^-8^), but no effect of coding model (*χ^2^*(2) = 3.7, *p* = .16). Importantly, there was an effect of the interactions coding model x type of outcome (*χ^2^*(4) = 54.5, *p* = 4.2 x 10^-11^, see Figure 4C) and type of outcome x ROI (*χ^2^*(16) = 31.9, *p* = .01), but no effect of the interactions ROI x coding model (*χ^2^*(16) = 3.0, *p* = ∼1) nor coding model x outcome type x ROI (*χ^2^*(32) = 16.2, *p* = .99). Post hoc tests (see “Univariate and multivariate effects of outcome valence”) showed that the absolute model had higher prediction accuracy than both the partially-adaptive (*p* = .016) and the fully-adaptive (*p* < .0001) models for factual outcomes with partial feedback, while for both factual and counterfactual outcomes with complete feedback the fully-adaptive model was more accurate than the absolute model (*p* = .001 and *p* = .004, respectively) and comparable to the partially-adaptive one (*p* = .35 and *p* = .19, respectively). Finally, there was no significant difference in prediction accuracy between the three outcome conditions in the most ventral region (all *p*s > .05), while all pairwise comparisons between outcome conditions were significant in frontopolar cortex (all *p*s < .03); in the other regions prediction accuracy was higher for factual than for counterfactual outcomes (all *p*s < .011) but it was similar for the two factual outcome types (all *p*s > .08).

To sum up, a formal comparison of three different coding models shows that outcome information encoding is more in line with absolute coding when only partial information about the outcome is provided but with fully-adaptive coding when counterfactual information is also presented. This suggests that making information about the alternative outcome available changes how the brain represents valence. We found poor prediction accuracy for all models in the most anterior region of mPFC. This could suggest that this area may encode outcome valence yet in a different way. However, we refrain from providing interpretations of differences between different ROI as accuracy measures depend on additional factors beyond neural encoding which usually differ between PFC regions (see Etzel et al., 2013; Bhandari et al., 2018 for a thorough discussion).

## DISCUSSION

We systematically investigated outcome processing and encoding in multiple subdivisions of mPFC/OFC and cingulate cortex, using different analysis approaches, to assess (1) univariate and multivariate effects in these regions, (2) possible effects of feedback information on outcome encoding, and (3) to compare different coding models of outcome valence.

A first goal of this study was to assess univariate and multivariate effects of outcome valence and feedback information in different regions in mPFC/OFC and cingulate cortex. We showed that signal change reflecting differences between good and bad outcomes was higher for factual than for counterfactual outcomes with no difference between the two factual outcomes (with complete or partial feedback) in aOFC, FPC, and vACC; in dSFG and mSFG the effect was higher for counterfactual than for factual outcomes, and in the remaining regions, there was no difference between the three outcome types. Additionally, while factual outcome processing generally increased activation in anterior ROIs (except pOFC), the same regions were deactivated by counterfactual outcome processing. Deactivation for counterfactual outcome processing in OFC replicates results from a previous study showing enhanced OFC activity for counterfactual processing in loss trials and relative deactivation in win trials in the complete feedback condition (Coricelli et al., 2005). Here, we did not distinguish between win and loss trials; however, as participants were highly accurate in their choices (see Figure 2), winning trials were much more frequent, thus, on average, deactivation should prevail, as we actually observed. As for the multivariate effects, we found that multivariate decoding was higher for factual than for counterfactual outcomes. Nevertheless, decoding accuracy for counterfactual outcomes was significantly higher than chance level, indicating that mPFC regions did encode counterfactual information. As counterfactual outcomes did not affect participants’ actual gains/losses (i.e., they were irrelevant for performance) and it has been shown that decoding accuracy is higher for attended than unattended (i.e., behaviorally irrelevant) items (e.g., Christophel et al., 2018), it is possible that counterfactual outcomes received less attention. Although univariate and multivariate results look slightly different, these differences are not rare (e.g., Mohr et al., 2015; Baggio et al., 2016; see Davis et al., 2014 for a thorough discussion of their interpretation). Notwithstanding, all regions that were selected showed univariate/multivariate effects, thus confirming their role in processing/encoding of outcome valence.

The second aim of the study was to investigate how outcome valence is represented in the brain. To this purpose, we compared the accuracy of three different coding models (i.e., absolute, partially-adaptive, and fully-adaptive) in predicting valence encoding in the different outcome conditions. We found that outcome valence is represented with different neural codes when either complete or partial feedback is provided. In particular, while neural coding is better described by absolute coding when partial feedback is given, relative coding fits the fMRI data better when feedback is complete. This result is in line with previous research using computational modeling and showing a better fit of behavioral data for an absolute coding model when only partial feedback is used, while a relative model fits the data better when complete feedback trials are also introduced (Bavard et al., 2018). In addition, our results are in line with previous research providing evidence for absolute coding of value signals in posterior ventromedial PFC when feedback information is partial (Burke et al., 2016). Interestingly, this evidence comes from univariate analyses, while the same study reports evidence for partially-adaptive coding from multivariate results. Burke et al. (2016) assessed different coding models by using two procedures: cross-set classification to test for partially-adaptive/absolute and fully-adaptive coding of value and support vector regression to adjudicate between partially-adaptive and absolute coding. The first method provides a measure of classification accuracy, while the second estimates prediction accuracy of the model, which are different measures, both qualitatively and conceptually. The difference between univariate and multivariate results may thus be due to comparing heterogeneous measures in the latter analysis. By using the same measure for all models, we showed that actually an absolute coding model fits the data better than a partially-adaptive one when feedback is partial.

The final goal of our research was to clarify how counterfactual information is represented and processed and to assess whether and how it affects outcome valence coding. First, we were able to decode counterfactual information from local activity patterns in the brain regions we considered. Moreover, we showed that counterfactual information processing reflects into deactivation/activation of the regions in the network, with an opposite pattern compared to factual information. Thus, counterfactual outcomes are both represented and processed in these regions. Our finding that in the most ventral mPFC areas factual outcomes are encoded positively and counterfactual outcomes are encoded negatively is consistent with previous evidence (Li and Daw, 2011; Klein et al., 2017). We extend these results by showing that the pattern is opposite in the dorsal mPFC areas. Finally, we found that presenting information about the alternative choice outcome affects how the brain encodes outcome valence, moving from an absolute code when information is partial to a relative code when information is complete, thus confirming our original hypothesis. But how does counterfactual information affect behavior? As previous studies showed that learning is significantly higher when complete information is provided than when feedback is partial (Palminteri et al., 2015, 2017; Bavard et al., 2018) and we found evidence for different coding of valence in distinct feedback conditions, the improvement in learning when complete feedback is given may arise from a more effective way (relative) of the brain to encode valence in this situation. Presenting information about the counterfactual outcome might make the context more salient, inducing a more effective representation of valence (relative instead of absolute). Given the structure of our task, in trials with complete feedback participants often experienced trials where factual and counterfactuals outcomes differed. This difference likely prompted a comparison between the two outcomes, making the value range more explicit and possibly inducing coding in terms of context. Although participants learned the associations between symbols and outcomes (they started choosing the best symbols more often after learning, see Palminteri et al., 2015) and could in principle anticipate the counterfactual outcome, they did not seem to represent it. We found evidence for absolute coding in trials with partial feedback; this result is compatible with the counterfactual outcome not being represented in this condition, indicating that a direct comparison between factual and counterfactual outcomes is crucial to represent valence in relative terms. Importantly, relative coding of valence is not only efficient, but it also produces behavioral effects: through normalization, neutral outcomes in the punishment context acquire positive value, thus punishment avoidance acts as reinforcement for choosing the symbol leading more likely to this outcome. Indeed, we find better performance in complete information trials when rescaling should occur (see Figure 2). This process induces symmetry between reward and punishment learning and can account for asymmetry effects previously reported with partial feedback (Kubanek et al., 2015). Our results provide evidence for this neural mechanism that was just hypothesized by testing different learning models (Palminteri et al., 2015). Although our neural results may also be explained by assuming that participants implement a Q-learning model in the partial feedback condition and a policy gradient (or direct comparison) model in the complete feedback condition (Li and Daw, 2011; Klein et al., 2017), these alternative models are not able to explain the behavioral patterns of preferences observed in previous studies (Palminteri et al., 2015; Bavard et al., 2018).

To summarize, this study offers the first systematic investigation of both univariate and multivariate effects of different information conditions on outcome valence encoding within multiple regions of mPFC/OFC and cingulate cortex. In addition, we provide the first comparison of three possible coding models (absolute, partially-adaptive, and fully-adaptive) for outcome valence encoding in the human brain by using comparable measures (i.e., obtained using the same analysis approach) of model accuracy in predicting the observed fMRI data. Taken together, our results help build a more comprehensive picture of how the human brain encodes and processes outcome valence. In particular, our results suggest that outcome valence is represented through multiple coding mechanisms, flexibly activated depending on the specific choice setting. We acknowledge some limitations of our study: first, in our experiment, we showed factual and counterfactual outcomes together, so activity related to either of them is not temporally separated. Moreover, different from previous studies investigating the pure representation of other cognitive variables (e.g., Reverberi et al., 2012; Pischedda et al., 2017), we investigated valence representation during outcome receipt, when additional cognitive processes (e.g., comparison between actual and expected outcome) likely took place. It should be noted, however, that it is extremely hard to dissociate outcome representation from the related computations (e.g., prediction error calculation) as these are triggered upon outcome presentation. Nonetheless, we hope our findings will stimulate future research to fully understand value-based decision, eventually.

## ACKNOWLEDGMENTS

This research was supported by a European Research Council Consolidator Grant “Transfer Learning within and between brains” (TRANSFER-LEARNING; agreement No. 617629). D.P. is currently supported by the Deutsche Forschungsgemeinschaft (DFG, German Research Foundation) under Germany’s Excellence Strategy “Science of Intelligence” (EXC 2002/1; project number 390523135). S.P. is supported by an ATIP-Avenir grant (R16069JS), the Programme Emergence(s) de la Ville de Paris, the Fyssen Foundation and the Fondation Schlumberger pour l’Education et la Recherche (FSER).

We thank Kai Görgen for his help in preparing the permutation test scripts.

## CONFLICT OF INTEREST

The authors declare no competing financial interests.

## AUTHOR CONTRIBUTIONS

D.P., S.P., and G.C. designed research; D.P. analyzed data; S.P. and G.C. provided input on data analysis; D.P. wrote the first draft of the paper; D.P. wrote the paper; D.P., S.P., and G.C. edited the paper.

## REFERENCES

1. Baggio G, Cherubini P, Pischedda D, Blumenthal A, Haynes J-D, Reverberi C (2016) Multiple neural representations of elementary logical connectives. NeuroImage 135:300–310.

2. Barr DJ, Levy R, Scheepers C, Tily HJ (2013) Random effects structure for confirmatory hypothesis testing: Keep it maximal. J Mem Lang 68:255–278.

3. Bates D, Mächler M, Bolker B, Walker S (2015) Fitting Linear Mixed-Effects Models Using lme4. J Stat Softw 67:1–48.

4. Bavard S, Lebreton M, Khamassi M, Coricelli G, Palminteri S (2018) Reference-point centering and range-adaptation enhance human reinforcement learning at the cost of irrational preferences. Nat Commun 9:4503.

5. Bermudez MA, Schultz W (2010) Reward Magnitude Coding in Primate Amygdala Neurons. J Neurophysiol 104:3424–3432.

6. Bhandari A, Gagne C, Badre D (2018) Just above Chance: Is It Harder to Decode Information from Prefrontal Cortex Hemodynamic Activity Patterns? J Cogn Neurosci 30:1473–1498.

7. Bunzeck N, Dayan P, Dolan RJ, Duzel E (2010) A common mechanism for adaptive scaling of reward and novelty. Hum Brain Mapp 31:1380–1394.

8. Burke CJ, Baddeley M, Tobler PN, Schultz W (2016) Partial Adaptation of Obtained and Observed Value Signals Preserves Information about Gains and Losses. J Neurosci 36:10016–10025.

9. Carandini M, Heeger DJ (2012) Normalization as a canonical neural computation. Nat Rev Neurosci 13:51–62.

10. Christophel TB, Iamshchinina P, Yan C, Allefeld C, Haynes J-D (2018) Cortical specialization for attended versus unattended working memory. Nat Neurosci 21:494–496.

11. Clithero JA, Carter RM, Huettel SA (2009) Local pattern classification differentiates processes of economic valuation. NeuroImage 45:1329–1338.

12. Clithero JA, Rangel A (2014) Informatic parcellation of the network involved in the computation of subjective value. Soc Cogn Affect Neurosci 9:1289–1302.

13. Conen KE, Padoa-Schioppa C (2019) Partial Adaptation to the Value Range in the Macaque Orbitofrontal Cortex. J Neurosci 39:3498 –3513.

14. Coricelli G, Critchley HD, Joffily M, O’Doherty JP, Sirigu A, Dolan RJ (2005) Regret and its avoidance: a neuroimaging study of choice behavior. Nat Neurosci 8:1255–1262.

15. Cox DD, Savoy RL (2003) Functional magnetic resonance imaging (fMRI) “brain reading”: detecting and classifying distributed patterns of fMRI activity in human visual cortex. NeuroImage 19:261–270.

16. Cox KM, Kable JW (2014) BOLD Subjective Value Signals Exhibit Robust Range Adaptation. J Neurosci 34:16533–16543.

17. Davis T, LaRocque KF, Mumford JA, Norman KA, Wagner AD, Poldrack RA (2014) What do differences between multi-voxel and univariate analysis mean? How subject-, voxel-, and trial-level variance impact fMRI analysis. NeuroImage 97:271–283.

18. Diekhof EK, Kaps L, Falkai P, Gruber O (2012) The role of the human ventral striatum and the medial orbitofrontal cortex in the representation of reward magnitude – An activation likelihood estimation meta-analysis of neuroimaging studies of passive reward expectancy and outcome processing. Neuropsychologia 50:1252–1266.

19. Duda RO, Hart PE, Stork DG (2000) Pattern classification, 2nd Ed. New York, NY, US: John Wiley & Sons.

20. Elliott R, Agnew Z, Deakin JFW (2008) Medial orbitofrontal cortex codes relative rather than absolute value of financial rewards in humans: Medial OFC and relative value. Eur J Neurosci 27:2213–2218.

21. Etzel JA, Zacks JM, Braver TS (2013) Searchlight analysis: Promise, pitfalls, and potential. NeuroImage 78:261–269.

22. Fan L, Li H, Zhuo J, Zhang Y, Wang J, Chen L, Yang Z, Chu C, Xie S, Laird AR, Fox PT, Eickhoff SB, Yu C, Jiang T (2016) The Human Brainnetome Atlas: A New Brain Atlas Based on Connectional Architecture. Cereb Cortex 26:3508–3526.

23. Friston KJ, Glaser DE, Henson RNA, Kiebel S, Phillips C, Ashburner J (2002) Classical and Bayesian Inference in Neuroimaging: Applications. NeuroImage 16:484–512.

24. Hare TA, O’Doherty J, Camerer CF, Schultz W, Rangel A (2008) Dissociating the Role of the Orbitofrontal Cortex and the Striatum in the Computation of Goal Values and Prediction Errors. J Neurosci 28:5623–5630.

25. Haxby JV, Gobbini MI, Furey ML, Ishai A, Schouten JL, Pietrini P (2001) Distributed and Overlapping Representations of Faces and Objects in Ventral Temporal Cortex. Science 293:2425–2430.

26. Hebart MN, Kai Görgen, Haynes J-D (2015) The Decoding Toolbox (TDT): a versatile software package for multivariate analyses of functional imaging data. Front Neuroinformatics 8:88.

27. Holm S (1979) A Simple Sequentially Rejective Multiple Test Procedure. J Stat 6:65–70.

28. Howard JD, Kahnt T, Gottfried JA (2016) Converging prefrontal pathways support associative and perceptual features of conditioned stimuli. Nat Commun 7:11546.

29. Kahnt T (2018) A decade of decoding reward-related fMRI signals and where we go from here. NeuroImage 180:324–333.

30. Kahnt T, Heinzle J, Park SQ, Haynes J---D (2010) The neural code of reward anticipation in human orbitofrontal cortex. Proc Natl Acad Sci 107:6010–6015.

31. Kahnt T, Heinzle J, Park SQ, Haynes JD (2011) Decoding different roles for vmPFC and dlPFC in multi-attribute decision making. NeuroImage 56:709–715.

32. Kahnt T, Park SQ, Haynes JD, Tobler PN (2014) Disentangling neural representations of value and salience in the human brain. Proc Natl Acad Sci 111:5000–5005.

33. Kennerley SW, Wallis JD (2009) Evaluating choices by single neurons in the frontal lobe: outcome value encoded across multiple decision variables. Eur J Neurosci 29:2061– 2073.

34. Klein TA, Ullsperger M, Jocham G (2017) Learning relative values in the striatum induces violations of normative decision making. Nat Commun 8:16033.

35. Knutson B, Fong GW, Adams CM, Varner JL, Hommer D (2001) Dissociation of reward anticipation and outcome with event-related fMRI. Neuroreport 12:3683–3687.

36. Knutson B, Fong GW, Bennett SM, Adams CM, Hommer D (2003) A region of mesial prefrontal cortex tracks monetarily rewarding outcomes: characterization with rapid event-related fMRI. NeuroImage 18:263–272.

37. Kobayashi S, Pinto de Carvalho O, Schultz W (2010) Adaptation of Reward Sensitivity in Orbitofrontal Neurons. J Neurosci 30:534–544.

38. Kriegeskorte N, Simmons WK, Bellgowan PSF, Baker CI (2009) Circular analysis in systems neuroscience: the dangers of double dipping. Nat Neurosci 12:535–540.

39. Kubanek J, Snyder LH, Abrams RA (2015) Reward and punishment act as distinct factors in guiding behavior. Cognition 139:154–167.

40. Lenth R (2018) emmeans: Estimated Marginal Means, aka Least-Squares Means. R Package version 1.1.3. https://CRAN.R-project.org/package=emmeans.

41. Li J, Daw ND (2011) Signals in Human Striatum Are Appropriate for Policy Update Rather than Value Prediction. J Neurosci 31:5504–5511.

42. Louie K, De Martino B (2014) The Neurobiology of Context-Dependent Valuation and Choice. In: Neuroeconomics, 2nd ed., pp 455–476. London, UK: Academic Press.

43. Louie K, Glimcher PW (2012) Efficient coding and the neural representation of value. Ann N Y Acad Sci 1251:13–32.

44. Mitchell TM (1997) Machine Learning. New York, NY, US: McGraw-Hill.

45. Mohr H, Wolfensteller U, Frimmel S, Ruge H (2015) Sparse regularization techniques provide novel insights into outcome integration processes. NeuroImage 104:163– 176.

46. Müller KR, Mika S, Ratsch G, Tsuda K, Scholkopf B (2001) An introduction to kernel-based learning algorithms. IEEE Trans Neural Netw 12:181–201.

47. Nieuwenhuis S, Heslenfeld DJ, Alting von Geusau NJ, Mars RB, Holroyd CB, Yeung N (2005) Activity in human reward-sensitive brain areas is strongly context dependent. NeuroImage 25:1302–1309.

48. Oldham S, Murawski C, Fornito A, Youssef G, Yücel M, Lorenzetti V (2018) The anticipation and outcome phases of reward and loss processing: A neuroimaging meta-analysis of the monetary incentive delay task. Hum Brain Mapp 39:3398–3418.

49. Padoa-Schioppa C (2009) Range-Adapting Representation of Economic Value in the Orbitofrontal Cortex. J Neurosci 29:14004–14014.

50. Padoa-Schioppa C, Assad JA (2006) Neurons in the orbitofrontal cortex encode economic value. Nature 441:223–226.

51. Palminteri S, Khamassi M, Joffily M, Coricelli G (2015) Contextual modulation of value signals in reward and punishment learning. Nat Commun 6:8096.

52. Palminteri S, Lefebvre G, Kilford EJ, Blakemore SJ (2017) Confirmation bias in human reinforcement learning: Evidence from counterfactual feedback processing. PLOS Comput Biol 13:e1005684.

53. Pischedda D, Görgen K, Haynes JD, Reverberi C (2017) Neural Representations of Hierarchical Rule Sets: The Human Control System Represents Rules Irrespective of the Hierarchical Level to Which They Belong. J Neurosci 37:12281–12296.

54. Rangel A, Camerer C, Montague PR (2008) A framework for studying the neurobiology of value-based decision making. Nat Rev Neurosci 9:545–556.

55. Reverberi C, Görgen K, Haynes JD (2012) Compositionality of Rule Representations in Human Prefrontal Cortex. Cereb Cortex 22:1237–1246.

56. Rohe T, Weber B, Fliessbach K (2012) Dissociation of BOLD responses to reward prediction errors and reward receipt by a model comparison: Dissociation of reward prediction errors and reward receipt. Eur J Neurosci 36:2376–2382.

57. Rustichini A, Conen KE, Cai X, Padoa-Schioppa C (2017) Optimal coding and neuronal adaptation in economic decisions. Nat Commun 8:1208.

58. Schoenbaum G, Saddoris MP, Stalnaker TA (2007) Reconciling the Roles of Orbitofrontal Cortex in Reversal Learning and the Encoding of Outcome Expectancies. Ann N Y Acad Sci 1121:320–335.

59. Stott JJ, Redish AD (2014) A functional difference in information processing between orbitofrontal cortex and ventral striatum during decision-making behaviour. Philos Trans R Soc B Biol Sci 369:20130472–20130472.

60. Tukey JW (1949) Comparing Individual Means in the Analysis of Variance. Biometrics 5:99–114.

61. Vickery TJ, Chun MM, Lee D (2011) Ubiquity and Specificity of Reinforcement Signals throughout the Human Brain. Neuron 72:166–177.

62. Wisniewski D, Reverberi C, Momennejad I, Kahnt T, Haynes JD (2015) The Role of the Parietal Cortex in the Representation of Task-Reward Associations. J Neurosci 35:12355–12365.

63. Worsley KJ, Friston KJ (1995) Analysis of fMRI Time-Series Revisited—Again. NeuroImage 2:173–181.

64. Yan C, Su L, Wang Y, Xu T, Yin D, Fan M, Deng C, Hu Y, Wang Z, Cheung EFC, Lim KO, Chan RCK (2016) Multivariate Neural Representations of Value during Reward Anticipation and Consummation in the Human Orbitofrontal Cortex. Sci Rep 6:29079.

